# Suspect screening-data independent analysis workflow for the identification of arsenolipids in marine standard reference materials

**DOI:** 10.1101/2024.08.31.610588

**Authors:** Shubhra Bhattacharjee, Miguel A. Chacon-Teran, Michael Findlater, Stacey M. Louie, Jeremy D. Bailoo, Amrika Deonarine

## Abstract

There has been limited research into arsenolipid toxicological risks and health-related outcomes due to challenges with their separation, identification, and quantification within complex biological matrices (e.g., fish, seaweed). Analytical approaches for arsenolipid identification such as suspect screening have not been well documented and there are no certified standard reference materials, leading to issues with reproducibility and uncertainty regarding the accuracy of results. In this study, a detailed workflow for the identification of arsenolipids utilizing suspect screening coupled with data independent analysis is presented and applied to three commercially available standard reference materials (Hijiki seaweed, dogfish liver, and tuna). Hexane and dichloromethane/methanol extraction, followed by reversed-phase high-performance liquid chromatography-inductively coupled plasma mass spectrometry and liquid chromatography-electrospray ionization-quadrupole time-of-flight mass spectrometry. Using the workflow developed, mass fragmentation matching, mass error calculations, and retention time matching were performed to identify suspect arsenolipids. Arseno-fatty acids (AsFAs), arsenohydrocarbons (AsHCs), and arsenosugar phospholipids (AsSugPLs) were identified with high confidence; AsHC332, AsHC360, and AsSugPL720 in seaweed, AsHC332 in tuna, and AsFA474 and AsFA502 in the dogfish liver. AsHC332, AsHC360, and AsFA502 were identified as promising candidates for further work on synthesis, quantification using MS/MS, and toxicity testing.

## 1.0 Introduction

Arsenolipids constitute a group of lipid-soluble organoarsenicals that have generated interest in recent years due to their occurrence in seaweed, fish oil, and fish. Arsenolipids occur as numerous chemical species and so far include: hydrocarbons (AsHC)^1-9^, fatty acids (AsFA)^1, 3, 5, 7, 9-11^, fatty alcohols^9, 12,^ phosphatidylethanolamine (AsPE)^13, 14^, triacylglycerols (AsTAG)^15^, phosphatidylcholine (AsPC)^14, 15^, phosphatidylinositols (AsPI)^15^, phosphatidylglycerols (AsPG)^15^, and arsenosugar phospholipids (AsSugar-PLs)^4, 9^). There is limited data on toxicological effects of arsenolipids^16-18^, though the evidence thus far suggests that there is the potential for toxic effects to humans. *In vivo* and *in vitro* testing have been performed on a single AsFA, three AsHCs, and more recently, on an AsTAG and AsPC ^17, 19-22^. AsHC332, AsHC360, and AsHC444 are cytotoxic to liver^17, 21^, bladder^17, 21^, epithelial^20^, and astrocyte cells^23^. There are some limitations to these studies - there is such little data on the occurrence and concentrations of arsenolipid compounds in seaweed and fish, that the few arsenolipids for which toxicity data is available may not be prevalent in seafoods and the exposure concentrations tested may not be relevant to human exposure.

Concentrations of total arsenolipids in seaweed and fish in the range 10^0^ - 10^2^ ng/g^24^ and up to 10 mg/kg in fish oil^25^ have been reported, but the majority of this data is based on the proportion of total arsenic that is present as lipid soluble or extractable in non-polar solvents, rather than concentrations of individual arsenolipids. A few studies have estimated individual arsenolipid concentrations by LC-ICPMS but good separation and subsequently accurate quantification of individual arsenolipid compounds using published methods is difficult to achieve. Often, arsenolipids have quantified as groups of compounds (e.g., sum of all AsFAs, AsHC332+AsHC360+AsHC404)^25, 26^. Individual concentration data for only two arsenolipids, As FA362 and AsHC360, for which synthesis methods exist, have been documented in algae, fish, and shellfish (33 - 40 ng/g) in Japan^25^.

Arsenic toxicity is a function of its speciation, and so to adequately address the question of arsenolipid exposure and toxicity, it is important to first determine the prevalence and relevant exposure concentrations of individual arsenolipids in seafoods. Arsenolipid identification and quantification in biological matrices has progressed from the use of liquid chromatography-inductively coupled plasma mass spectrometry (LC-ICPMS), where arsenolipid peaks are detected as arsenic (*m/z* 75), to coupling LC-ICPMS with electrospray ionization (ESI) high-resolution mass spectrometry (HRMS) for the molecular identification of arsenolipids through non-targeted and suspect screening approaches^27-29^. The use of molecular identification using ESI is necessary due to species co-elution and the destructive nature of the ICP, which is problematic for the separation and accurate identification of compounds^30^. Suspect screening has often been preferred for compound identification to avoid the limitations of non-targeted analysis in determining chemical structures accurately^31^. Even so, the identification of discrete arsenolipids with a high degree of confidence remains challenging. One reason for this is difficulties in replicating suspect screening methods due to limited documentation of the MS data processing. Most studies inadequately describe workflow details such as the full selection of precursor ions utilized, the mass error tolerance for matching of precursor and fragmented masses, and screening criteria for the selection of LC peaks for analysis (e.g., signal to noise ratio, retention time matching tolerance). These parameters are crucial for analysis and ensuring good data integrity; for example, increasing the mass error tolerance from 1 to 10 ppm or the exclusion of a potential precursor ion can drastically affect the list of suspect compounds generated. Thus, there is a need for a highly detailed suspect screening data processing workflow for arsenolipid identification.

Furthermore, from an analytical perspective, arsenolipid standards for ensuring quality analysis/quality control are critically needed and will aid in improving data quality and the reproducibility of methods. Arsenolipids are not commercially available and are difficult to synthesize with high yield, even when synthesis methods exist^32^. Some researchers have used LC separation followed by fraction collection to obtain solutions of relatively high purity and concentration. This is time consuming and the potential for co-elution of arsenolipids makes collection of a high purity fraction difficult to achieve.

Standard reference materials (SRMs) or certified reference materials (CRMs) for arsenolipids would greatly assist in improving quality control and reliability of measurement, particularly as they are often easily purchased. However, there are currently no arsenolipid-certified SRMs. Seven to eight arsenolipids in the Hijiki seaweed CRM 7405-a were previously quantified by Glabonjat et al.^33^ and Amin et al.^25^ using LC-ICPMS/ESIMS but that SRM has since been discontinued and replaced by CRM 7405-b. Raber et al. later identified and quantified AsHC332 and AsHC360 in CRM 7405-b using LC-ICPMS and gas chromatography-MS/MS (GC-MS/MS)^34^; however, this CRM is not yet certified and there are likely multiple arsenolipids which remained unquantified.

The goals of this study were therefore to address the need for improved reporting and quality analysis/quality control of methods for arsenolipid identification. Specific objectives were to develop a suspect screening-data independent analysis workflow using LC-ICPMS and LC-ESI-QTOFMS and to apply it to three commercially available standard reference materials. Candidate arsenolipids for further work on synthesis, characterization, and LC-MS/MS or GC-MS/MS quantification were identified - a necessary first step towards arsenolipid-certified SRMs.

## 2.0. Materials and Methods

### 2.1 Reagents and Standards

Concentrated nitric acid (Optima grade, Fisher Scientific), hexane (Optima grade, Fisher Scientific), formic acid (Optima grade, Fisher Scientific), methanol (Optima grade, Fisher Scientific), and dichloromethane (HPLC grade, Alfa Aesar) were purchased. Arsenobetaine and dimethylarsinic acid standards (≥95% purity) were purchased from Sigma Aldrich. An arseno fatty acid, AsFA362, was synthesized using our previously established synthesis method^32^ and a stock solution of 1000 mg/L was prepared in methanol. Three standard reference materials were obtained: seaweed hijiki (CRM 7405-b) from the National Metrology Institute of Japan, dogfish liver (DOLT-5) from the National Resource Council Canada, and tuna fish (BCR-627) from the European Commission Joint Research Center. All reference materials were certified for total arsenic concentration^35,36^,37 (Table S1).

### 2.2 Arsenolipid extraction procedure

Varying organic solvents have been used for sample preparation in previous studies (e.g., hexane, dichloromethane/methanol, pyridine^9, 33, 38^). Here, arsenolipid sequential extraction was performed using an approach adapted from Petursdottir et al. (Figure S1) which employed a two step extraction using hexane followed by dichloromethane/methanol (DCM/MeOH)^9^. DCM/MeOH has been shown to work well for arsenolipid extraction^33^. The addition of a hexane extraction step was considered a reasonable approach to maximize the extraction of arsenolipids by using three organic solvents covering a range of log P values (hexane 2.59, DCM 1.19, MeOH -0.72^39^) ^9, 40^ and to facilitate comparison with previous studies which used different solvents but with log P values within this range. 0.5 g of sample was placed in a 50 mL polypropylene centrifuge tube and extracted with 4 mL hexane using a vertical rotating mixer (ATR Rotamix RKVSD, USA). The samples were shaken overnight (12 hr) at 50 rpm, at 21°C, and then left to stand for 10 minutes. The supernatant was then transferred to a 15 ml centrifuge tube, evaporated using a n-evap nitrogen evaporator (Organomation, USA) with ultra-high purity nitrogen gas and then reconstituted with 4 mL of MeOH. The residual fraction was dried using ultra high purity nitrogen gas, dispersed in 4 mL of DCM/MeOH (2:1, v/v), and then shaken overnight (12 hr) on a horizontal orbital shaker (Thermo Scientific MaxQ 4000 Benchtop, USA) at 200 rpm, at 21°C. The DCM/MeOH supernatant was then dried using ultra-high purity nitrogen gas and reconstituted in 4 mL of methanol. All samples were filtered (0.22 µm hydrophobic PTFE filter, MicroSolv) and stored at 4°C in the dark until LC-ICPMS and LC-ESI-QTOFMS analyses.

### 2.3. LC-ICPMS

Samples were analyzed using LC (Agilent 1260 Infinity II, USA) coupled with ICPMS (Agilent 7900, USA) to investigate the occurrence of lipid-soluble arsenic-containing compounds. The carrier gas was ultra high purity argon. A high purity 20% oxygen/80% argon gas mixture was used as a supplemental gas (15.5%) to minimize carbon deposition on the platinum cones and helium gas (4 mL/min) was used in the collision cell to minimize the formation of polyatomic spectral interferences with the same *m/z* as arsenic (*m/z* 75). A Poroshell 120 EC C18 reversed-phase column (4.6×100 mm, 2.7µm, Agilent, USA) was used with a gradient flow program. The mobile phase solvents were, A: ultrapure water with 0.05% (v/v) formic acid and, B: MeOH with 0.05% (v/v) formic acid. The gradient flow comprised the following steps: (i) 98% A/2% B (0 to 4 minutes); (ii) change from 98% A/2% B to 5% A/95% B (4 to 6 minutes); (iii) maintain 5% A/95% B (6 to 16 minutes); (iv) change from 5% A/95% B to 98% A/2% B (16 to 17 minutes); and (v) maintain 98% A/2% B (17 to 26 minutes). The injection volume and flow rate were 30 µL and 0.3 mL/min, respectively. Additional details about instrument settings are provided in Table S2.

Arsenolipids were quantified using a calibration curve prepared with dimethylarsinic acid calibration standards, which ranged from 0.1 to 10 µg/L. Only peaks that eluted between 6 and 17 minutes were considered for total arsenolipid quantification. The concentrations of all eluted arsenolipid peaks were summed and compared to the certified total arsenic concentration to calculate the percentage of total arsenic present as arsenolipids.

### 2.5. LC-ESI-QTOFMS

The reconstituted hexane and DCM/MeOH extracts were analyzed using LC (Agilent 1260 Infinity II, USA) coupled with ESI-QTOFMS (Agilent 6545 QTOF with DualJet AJS ESI, USA) to obtain high resolution mass spectra for identifying arsenolipids. The analysis utilized the same Poroshell 120 EC C18 reversed phase column, mobile phases, flow rate, and gradient flow program as the LC-ICPMS method. Mass spectra were collected in positive ion mode within the *m*/*z* 100 to 1700 range for full scan spectra and *m*/*z* 40 to 1700 for fragment spectra. A reference ion mixture (*m*/*z* 121.050873 and 922.009798) was constantly injected for reference mass correction. Source conditions were selected based on the ESI-QTOFMS settings reported by Al Amin and colleagues as follows: 100 °C drying gas temperature at 12 L/min flowrate, 45 psi nebulizer pressure, 350 °C sheath gas temperature at 11 L/min flowrate, 4500 V capillary voltage, 500 V nozzle voltage, and 135 V fragmentor voltage^25^. The data were collected using MassHunter Acquisition B.09.00. Additional information regarding the instrument settings is provided in Table S2. Two run modes were performed per sample: (1) all-ions mode, in which the QTOF repeatedly cycled between collecting a full scan TOFMS spectrum with no application of collision energy, followed by fragmentation of all ions (no precursor selection) at collision energies of 10, 20, and 40 V (referred to as MS Scan in Figure 1); and (2) auto-MS/MS in preferred ions only mode (referred to as MS/MS in Figure 1), in which precursor ions matching those in a database (described below) were selected in the quadrupole for fragmentation at collision energies of 10, 20, and 40 V and TOF analysis of the fragment ions. The preferred ions mode was implemented for MS/MS because preliminary runs without a preferred ion list generally failed to select any suspect arsenolipids from the database for fragmentation from the complex sample matrix. This method restricted MS/MS spectrum collection to only compounds in the database; however, the all-ions data set can potentially be data-mined to search for fragments of other suspect arsenolipid compounds of interest.

**Figure 1:**
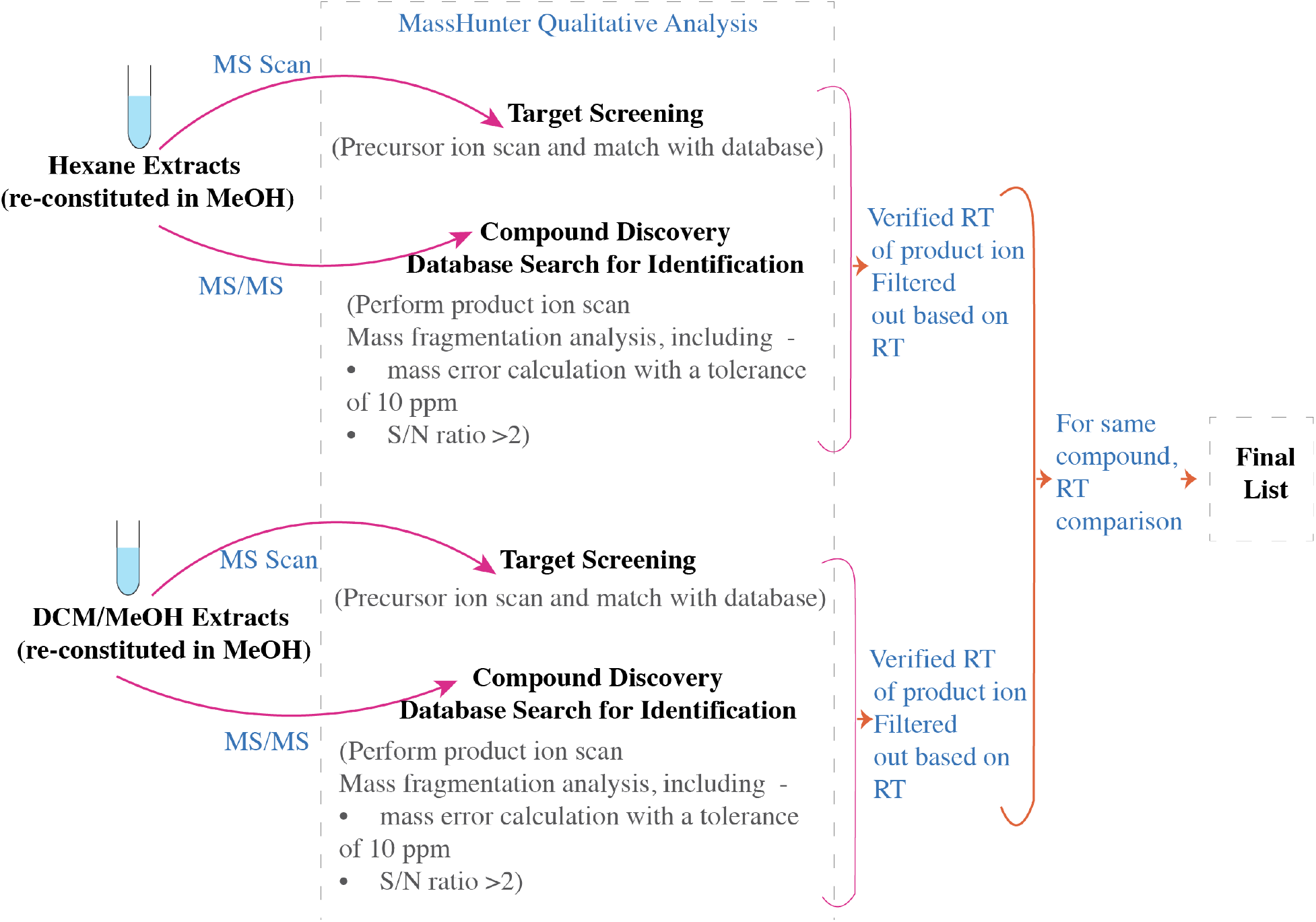
The workflow for suspect arsenolipids screening and data independent analysis. LC-ESI-QTOFMS was utilized to acquire high-resolution mass spectra for reconstituted hexane and DCM/MeOH extracts, which were subsequently analyzed using Mass Hunter Qualitative Analysis 10.0. Initial precursor ion screening and database matches were performed using the all ions run mode (MS Scan), followed by a similar procedure with precursor ion database matching and product ion filtering for expected arsenolipid fragments in auto MS/MS mode (MS/MS). Suspect arsenolipids were further filtered for mass error on the precursor and fragment masses, as well as signal to noise ratio of the fragment ion intensities. The retention times of MS/MS product ions were confirmed by comparing the chromatograms of precursor and product ions at collision energies of 0, 10, 20, and 40 V. Suspect arsenolipids identified in both MS Scan and MS/MS run modes were then compared based on retention time. Finally, if the same compound was identified in both hexane and DCM/MeOH fractions, retention times were further matched. Suspect compounds were identified with varying confidence levels based on the screening criteria listed.

### 2.5. Arsenolipids database

There was no publicly available arsenolipids database, so one was compiled using PCDL Manager B.08.00 (Table S3). A search of the Web of Science database was performed on 2024-05-08 using a title search with the keywords: “arsenolipids OR arsenic-containing fatty acid OR arsenic-containing hydrocarbons OR arsenolipid OR arsenohydrocarbon OR arsenofattyacid”. This search yielded 133 articles spanning from 1979-01-01 to 2024-03-23. Prior to 2008, there was limited information available on the masses and formulae of identified arsenolipids. Consequently, the search identified 102 articles published between 2008 and 2022, which was further narrowed to 30 articles with molecular formulas and mass information^1, 2, 4-9, 12-16, 19, 22, 25, 27, 29, 33, 38, 41-53^. The database included 154 arsenolipids identified within seafood and seaweed matrices (Table S3). This dataset included compound name, empirical formula, and exact mass of the neutral molecule. The masses of the [M+H]^+^ adducts for each arsenolipid are listed Table S3. Preliminary analyses of full scan MS data against the database revealed suspect compound matches for the sodium and ammonium adducts, as well as dehydration products. Therefore, to improve the comprehensiveness of the precursor selection, the *m*/*z* values for [M+H]^+^, [M+Na]^+^, [M+NH_4_]^+^, [M–H_2_O+H]^+^, [M–H_2_O+Na]^+^, and [M–H_2_O+NH_4_]^+^ were included for each compound in the preferred ion list of compounds to be fragmented in MS/MS mode.

### 2.6. Identification of arsenolipids

The high-resolution mass spectra obtained using LC-ESI-QTOFMS were analyzed and processed using MassHunter Qualitative Analysis 10.0. The analysis workflow (Figure 1) involved first performing target screening against the database on the MS data, allowing H^+^, Na^+^, and NH_4_ ^+^ for the positive ion charge carriers, as well as neutral losses of H_2_O. The mass error tolerance for the target screening was 10 ppm. Mass errors were also manually calculated on the extracted suspect compounds to ensure that only those with a mass error ≤ 10 ppm were accepted for further evaluation.

The MS/MS data were then processed using a MassHunter Compound Discovery search with fragment mass filtering to extract compounds having any of the following mass fragments: *m*/*z* 102.9529, 104.9685, 122.9791, 237.0103, or 409.0244 (mass match tolerance of ± 0.5 *m*/*z*). These masses have been previously identified as common arsenolipid mass fragments (Table S4). The masses of the precursor ions for the extracted compounds were then matched against the PCDL database to generate a list of suspect arsenolipids, with the same criteria such as mass error tolerance, etc., applied as in the MS Scan database matching (i.e., charge carriers/neutral loss settings, mass error tolerance). Manual inspection of the fragment spectra was then conducted. Only those compounds with at least two of the three commonly reported fragments for arsenolipids (*m*/*z* 102.9529, 104.9685, and 122.9791) and with intensities more than twice the baseline were selected. Due to the limited available references on the mass fragmentation of AsSugPLs, a minimum of one fragment, either *m/z* 237.0103 or 409.0244, was considered for identifying suspect AsSugPLs^14^. Suspect compounds that met these criteria with mass error tolerances of ≤ 10 ppm for all fragment ions were assigned a high level of confidence. Suspect compounds with at least two matching mass fragments but having one fragment with mass error > ∼10 ppm were included in the analysis results with a lower confidence level. The retention times of the MS/MS product ions were validated by comparing the precursor ion chromatograms with product ion chromatograms at collision energies of 0, 10, 20, and 40 V in the MS Scan. Next, the suspect arsenolipids generated using the MS target screening and MS/MS compound discovery database search were compared based on retention time. Only suspect arsenolipids with similar retention times (±1 minute) in the two separate injections for each analysis run mode (MS Scan and MS/MS) were accepted. Finally, in cases where a compound was identified in both the hexane and DCM/MeOH extracts, only those with matching retention times (± 1 minute) in both extracts were retained in the suspect compound list.

## 3.0. Results and Discussion

### 3.1 Method validation

Method development was first performed using AsFA362 and arsenobetaine, another organoarsenical commonly detected in seafood that is commercially available. A 10 µg/L AsFA362 standard was prepared in methanol and a 10 µg/L arsenobetaine standard was prepared in water. The retention time for the AsFA362 standard was 13.6 minutes for LC-ICPMS (Figure 2), while for LC-ESI-QTOFMS the retention time was 13.03 minutes, and the precursor mass error was -3.31 ppm. The mass fragmentation pattern of AsFA362 indicated fragments with *m*/*z* 104.9678 (Δ*m* = 6.6 ppm) and 122.9778 (Δ*m* = 10.8 ppm), corresponding to the arsenic-containing fragments As(CH_3_)_2_^+^ and As(CH_3_)OH_2_ ^+^ (Figure 2). The observed fragmentation patterns were consistent with the earlier research findings of Liu and colleagues^27^ and Al Amin and colleagues^25^, who identified AsFA362 in marine fish. AsB was also successfully identified using the workflow. The retention time of AsB was compared and found to be consistent between LC-ICPMS (3.55 minutes) and LC-ESI-QTOF-MS (3.584 minutes) (Figure 2). The mass error was 5.61 ppm for the precursor ions, however, there was no mass fragmentation data for AsB.

**Figure 2:**
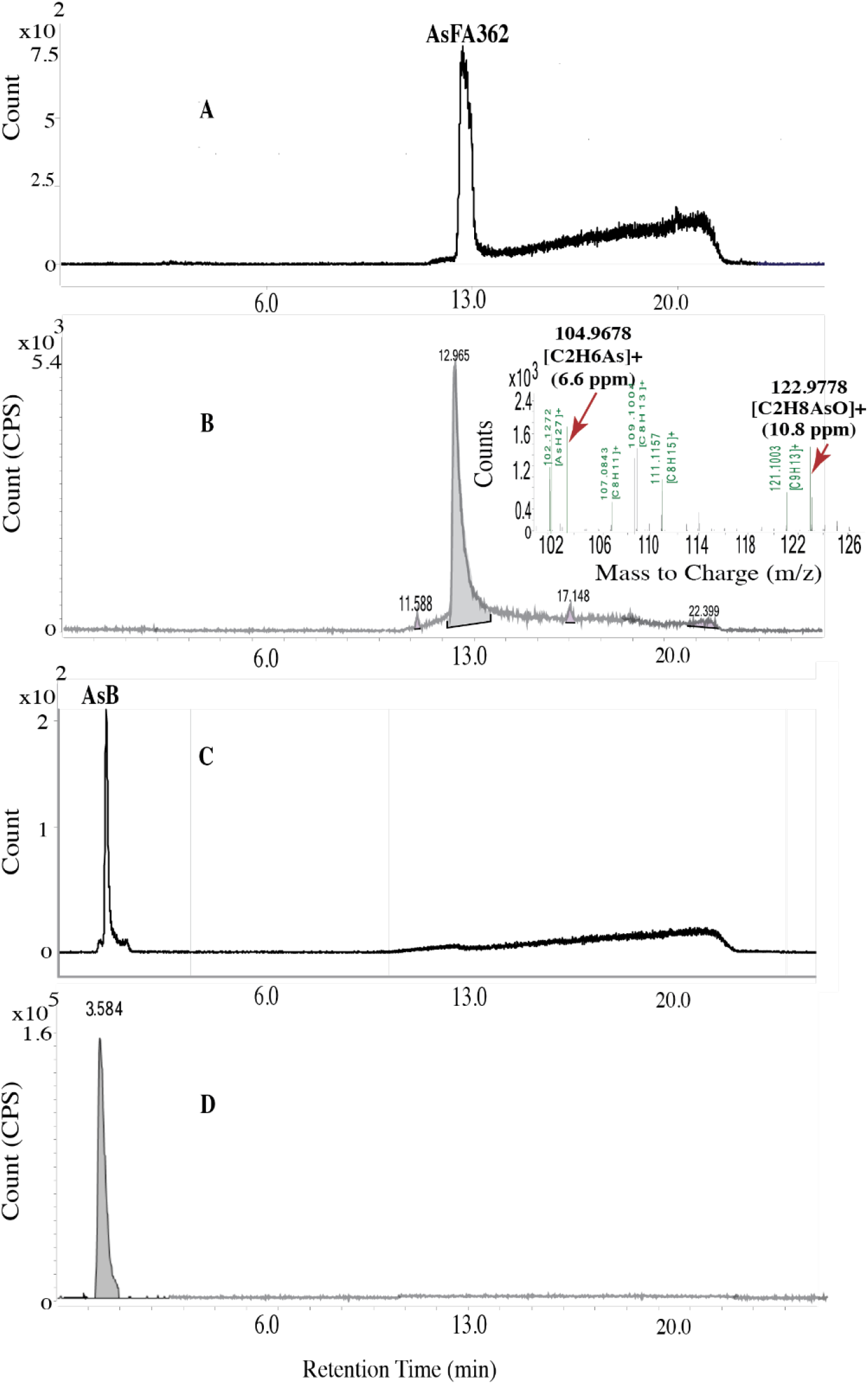
The chromatographic separation of AsFA362 was achieved using a Poroshell 120 EC C18 reverse-phase column (4.6×100 mm, 2.7µm) using LC-ICPMS (A). The extracted ion chromatogram of AsFA362 was obtained using reverse-phase LC-ESI-QTOFMS (B). The insert shows the MS/MS spectrum, which indicated the presence of *m*/*z* 104.9678 and 122.9778 fragments. The mass error was calculated in ppm for each fragment and is displayed within the parentheses. The chromatograms of AsB were obtained with LC-ICPMS (C) and LC-ESI-QTOFMS (D).

### 3.2. Arsenolipids in hijiki seaweed (NMIJ CRM 7405-b)

In the Hijiki seaweed SRM, arsenolipids ranged between 0.57 to 1.46% (504 ± 311 ng/g) of the total arsenic content (Table S6). Arsenolipids were found in both hexane and DCM/MeOH fractions; broad arsenolipid peaks were observed in the DCM/MeOH extract compared to the hexane extract (Figure 3). Six suspect arsenolipids with *m*/*z* values of 361.2469, 721.2926, 981.5068, 333.2147, 503.3463, and 749.3240 were identified, containing mass fragments *m*/*z* 104.9685, 122.9791, 237.0103, and/or 409.0244 (Figures 3, S2, S3). The arsenolipids meeting the high confidence criteria were: AsHC360 (precursor Δ*m* = 0.24 ppm), AsHC332 (precursor Δ*m* = -1.81 ppm), and AsSugPL720 (precursor Δ*m* = 0.55 ppm) (fragment mass errors are included in Figure 3). In the hexane fraction, AsHC360, AsSugPL720, and AsSugPL980 were detected, whereas AsSugPL720, AsSugPL748, AsFA502, and AsHC332 were identified in the DCM/MeOH fraction (Table S5). The current findings differ from previous results. AsSugPLs were detected in both hexane and DCM/MeOH, whereas earlier findings showed that AsSugPLs were more frequently identified in DCM/MeOH^33^. AsFA502 (precursor Δ*m* = -2.58 ppm), AsSugPL980 (precursor Δ*m* = -0.18 ppm), and AsSugPL748 (precursor Δ*m* = -3.06 ppm), were also identified but with a lower level of confidence (Figure S3). LC-ICPMS results also indicated additional arsenolipid peaks which remained unidentified. These could be arsenolipid species such as TMAsFOH, AsPE, and AsPC, however due to the lack of comprehensive mass fragmentation information for these compounds, they could not be identified using this workflow.

**Figure 3:**
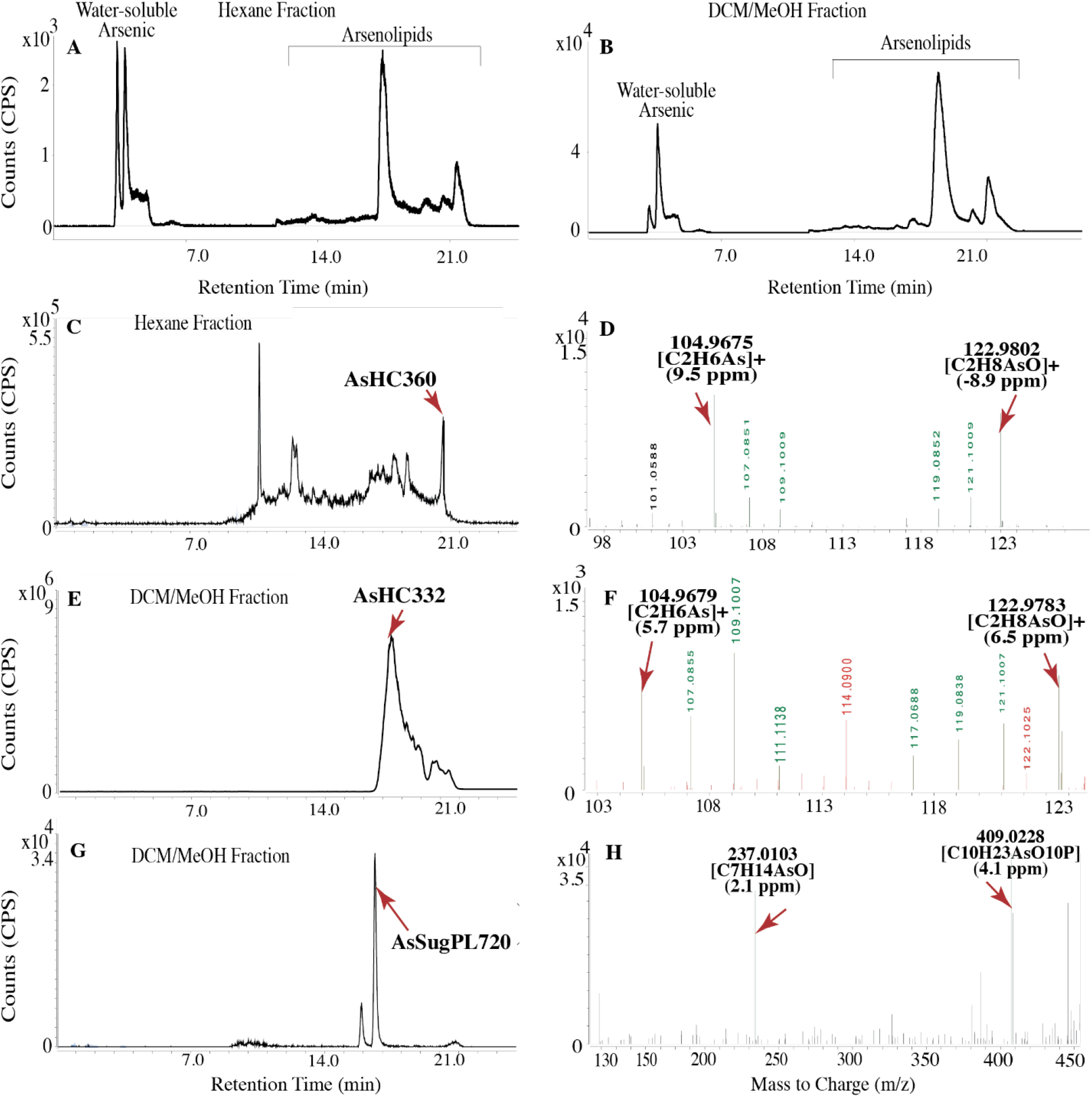
Arsenolipids in CRM 7405-b were separated chromatographically using LC-ICPMS (A and B) and identified using LC-ESI-QTOFMS (C, D, E, F, G, and H). In LC-ICPMS, detection was based on monitoring the *m*/*z* 75 signal. In LC-ESI-QTOFMS, the elution profiles of AsHC332, AsHC360, and AsSugPL720 were plotted as the extracted ion chromatogram (EIC) for the precursor ion obtained in the MS Scan (C, E, and G), with MS/MS sample runs used for mass fragment confirmation of arsenic-containing fragments and retention time matching on the EICs (F, D, and H). The mass error for the fragments is displayed within parentheses.

While AsHCs and AsSugPLs have often been the predominant arsenolipids found in seaweed^4^, this study reports for the first time the occurrence of an AsFA in seaweed (AsFA502). The identification of AsHC332 and AsHC360 in this study was consistent with previous work on hijiki seaweed CRM 7405-a and CRM 7405-b, in which AsHC332 and AsHC360 were also identified^33,34^. However, different AsSugPLs were previously identified in CRM 7405-a but were not identified in this study (e.g., AsSugPL958, AsSugPL986, AsSugPL1014, AsSugPL1042, AsSugPL1070)^33^, despite a similar extraction procedure using DCM/MeOH^33, 34^. Also, Glabonjat et al. used silica columns which can reduce arsenolipid detection limits^33^ but even so, there was no overlap with the AsSugPLs identified in this study. The absence of detailed information on fragmentation behavior, mass error calculation of product ions, and intensity considerations makes the verification of the AsSugPL results challenging.

### 3.3 Arsenolipids in dogfish liver (DOLT-5)

In the dogfish liver standard reference material, arsenolipids ranged between 0.004 to 0.005% (1.8 ± 0.4 ng/g) of the total arsenic content (Table S6). In the LC-ICPMS analysis, the DCM/MeOH extraction step yielded more arsenolipids from dogfish liver compared to the hexane extraction, indicating that DCM/MeOH was more effective in extracting arsenolipids from dogfish liver than hexane (Figure 4). Four arsenolipids with *m*/*z* values of 475.3163, 361.2450, 333.2130, and 503.3407 were identified with product ions with *m/z* 102.9529, 104.9685, and/or 122.9791 (Figures 4, S4, S5), corresponding to AsFA474 (precursor Δ*m* = -1.99 ppm), AsHC360 (precursor Δ*m* = 3.04 ppm), AsHC332 (precursor Δ*m* = 3.31 ppm), and AsFA502 (precursor Δ*m* = 0.19 ppm). AsFA474 and AsFA502 were identified with a high level of confidence (Figure 4), while AsHC332 and AsHC360 were identified with lower confidence (Figure S5). AsFA474 was identified in both the hexane and DCM/MeOH extracts (Table S5). AsHC360 was identified in the hexane extract, whereas AsHC332 and AsFA502 were identified in the DCM/MeOH extract. As with the hijiki seaweed SRM, additional peaks detected in LC-ICPMS could not be identified due to limited arsenolipid mass fragmentation data.

**Figure 4:**
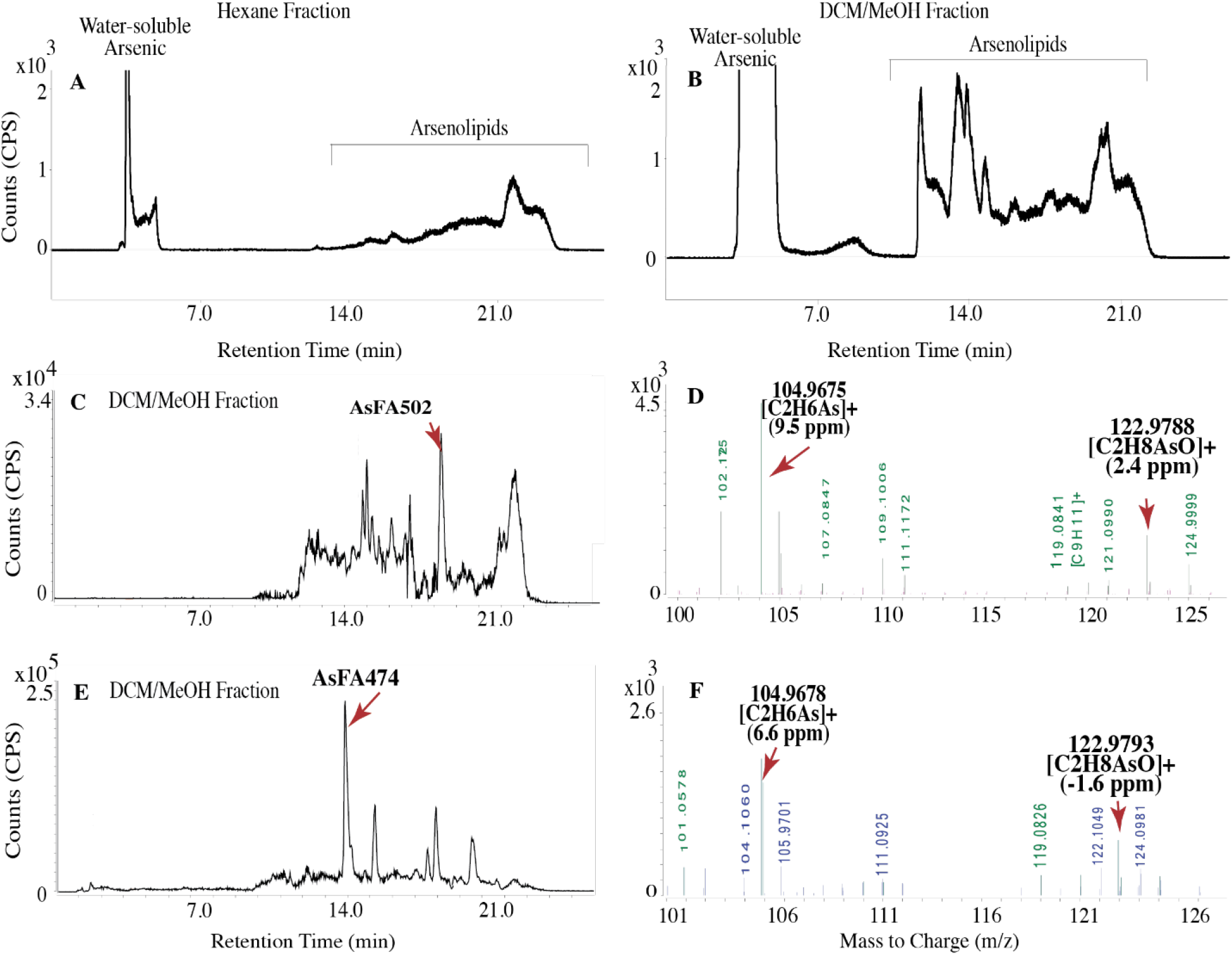
Arsenolipids in DOLT-5 were chromatographically separated using LC-ICPMS (A and B). The identification of AsFA502 and AsFA474 was accomplished with high confidence through LC-ESI-QTOFMS (C and E). The MS/MS spectra displayed mass fragments at m/z 102.9529, 104.9685, and 122.9791 (D, and F), with the fragment mass error indicated within the parentheses.

Previous research on arsenolipids exclusively employed dogfish liver for total arsenic analyses^16, 54^, neglecting investigations into arsenolipids, and therefore comparing findings was not possible.

However, the results of this study aligned with studies performed on other fish livers; AsHC332 and AsHC360 were identified in canned codfish liver tissue^2^ and in fresh codfish liver^3^. Sele and colleagues also identified AsHC332, AsHC360, and five AsFAs (AsFA362, AsFA388, AsFA390, AsFA436, and AsFA448) in another codfish liver study^16^. Based on current and previous findings, AsHC332 and AsHC360 are commonly present in fish livers. The other arsenolipids identified in codfish liver by Sele et al. were not identified in the dogfish liver SRM in this study, despite a similar sequential extraction procedure using hexane followed by MeOH^16^.

### 3.4. Arsenolipids in tuna fish (BCR-627)

In the tuna SRM, arsenolipids ranged between 0.002 to 0.009% (4.9 ± 0.5 ng/g) of the total arsenic content (Table S6). In LC-ESI-QTOFMS analysis, three suspect arsenolipids were identified with *m*/*z* values 333.2142, 359.2444, and 503.3484 (Figures 5, S6, S7), corresponding to AsHC332 (precursor Δ*m* = -0.31 ppm), AsHC358 (Δprecursor *m* = 1.91 ppm), and AsFA502 (precursor Δ*m* = -2.58 ppm). In the MS/MS spectra, product ions with *m/z* 102.9529, 104.9685, and/or 122.9791 were detected with mass errors ranging from 6.5 to 14.5 ppm. Only AsHC332 was identified with a high level of confidence (Figure 5). AsHC332 was identified in both the reconstituted hexane and DCM/MeOH extracts (Table S5), while AsFA502, and AsHC358 were identified only in DCM/MeOH extracts. For AsHC358 and AsFA502, the mass error calculation for *m/z* 104.9685 exceeded 10 ppm, which lowered the confidence in the identification of these compounds (Figure S7). AsHC332 has been previously identified in sashimi tuna^54^, skipjack tuna^38^, and tuna fish fillet ^29^. Other arsenolipids identified in tuna have included: AsHC404, AsHC406, AsHC360, AsFA362, AsFA390, AsFA348, AsFA418, AsFA376, AsFA404, AsFA448, AsFA528, and AsHC358^29, 38, 54^. However, these arsenolipids were not identified in this study. Solvents used for extraction were pyridine^38^, DCM/MeOH^29^, or chloroform/methanol^54^ which were either the same or within the log P range of solvents used in this study (log P: pyridine 0.73, chloroform 1.97^39, 55^), suggesting that sample extraction was not the critical issue, rather the uncertainty involved in processing of the MS data or simply sample heterogeneity.

**Figure 5:**
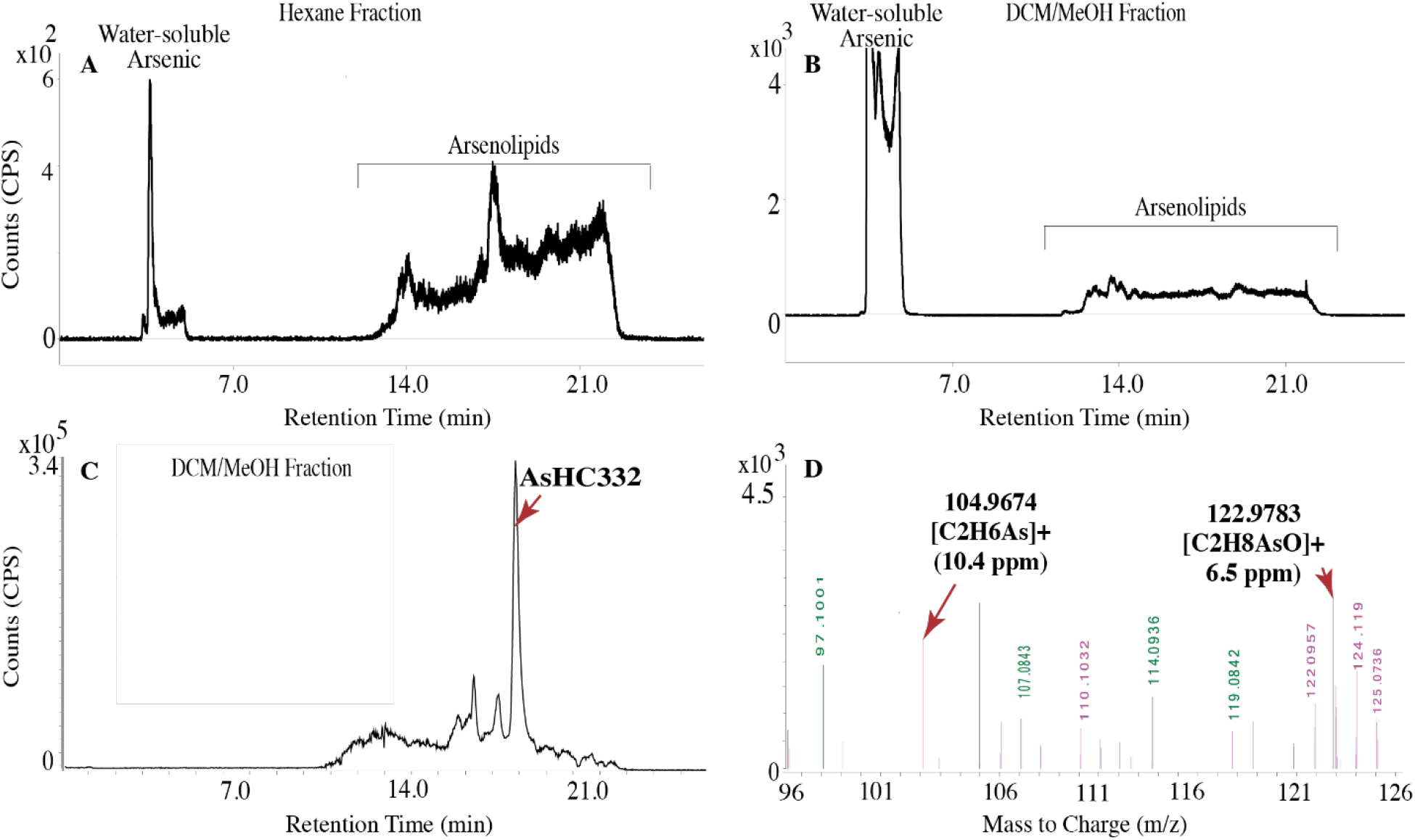
Chromatographic separation of arsenolipids in BCR-627 was performed using LC-ICPMS (A and B). MS/MS spectra confirmed the presence of AsHC332 with high confidence through LC-ESI-QTOFMS (C and D). The fragment mass error is shown within parentheses.

## 4.0 Conclusions and Future directions

This study is an initial step towards harmonizing arsenolipid identification approaches for marine biological matrices across studies, with highly detailed reporting of a suspect screening-data independent analysis workflow. Six arsenolipids in three marine SRMs were identified, within the sub-groups AsSugPLs, AsHCs, and AsFAs. Among the sub-groups of arsenolipids, AsSugPLs were exclusively found in the hijiki seaweed SRM (AsSugPL720), while only AsHCs and AsFAs were observed in the tuna (AsHC332) and dogfish liver (AsHC360, AsHC332) SRMs. These results corroborate previous work to some extent, despite differences in sample preparation steps, identification criteria, and instrument settings, which lends confidence to our results. With respect to solvents for extraction, our results suggested that an extraction step using a solvent with a higher log P compared to DCM/MeOH should be included, rather than the single step DCM/MeOH extraction that was proposed by Glabonjat et al.^33^, for example. The reason for this is that different arsenolipids were extracted with hexane versus DCM/MeOH in this study.

AsHC332 and AsFA502 were detected in all SRMs, albeit with varying levels of confidence, suggesting that they are appropriate candidates for further work on synthesis and the development of quantitative LC-MS analytical methods for these SRMs. Additionally, several potential arsenic-containing peaks with *m*/*z* 75 were not identified, indicating the occurrence of arsenolipids that were not listed in the PCDL database, underscoring the difficulty of definitively identifying relatively minor species such as TMAsFOH, AsPC, AsPI, etc., and the need for synthesis and characterization of these compounds, particularly mass fragmentation patterns and compound structure data.

Despite their discovery in 1979, there have been only a few toxicological studies on a limited set of compounds for which synthesis methods exist. A limiting factor here is that specific arsenolipids which are of concern in seafoods, due to high concentrations or wide prevalence, have not been identified. This is critical because the chemical speciation of arsenic present in the diet determines metabolism, biological uptake, toxicological endpoints and exposure risk. Our data suggests that AsHC332 and AsFA502 would be good candidates for toxicological testing, since they occurred in all three SRMs. AsHC360 is also a good candidate, as it was identified in the tuna SRM, and tuna is one of the third most widely consumed seafoods in the world^56^.

## Acknowledgements

The authors are grateful for the financial support received from the National Institute of Environmental Health Sciences (NIEHS) through grants R03ES033333 (A.D., J.B., M.F.) and R03ES034194 (A.D., J.B.), and the Graduate Student Research Program of Texas Tech University. The authors also thank Dr. Travis Falconer and Dr. Kevin Kubachka from the Food and Drug Administration who provided guidance on the identification of organoarsenicals in this study.

## Supplementary Information

The supplementary information includes data on total arsenic concentrations in SRMs, instrumental conditions for LC-ICPMS and LC-ESI-QTOFMS, a list of arsenolipids previously identified in marine biota and seafood SRMs, previously reported precursor ions and their mass fragments, and arsenolipids identified with high and low confidence in this study. Additionally, the supplementary figures illustrate the workflow of extraction, product ion chromatogram to compare retention times, MS/MS spectra, and extracted ion chromatograms of arsenolipids identified with lower confidence.

