## Supplementary Information for "Suspect screening-data independent analysis workflow for the identification of arsenolipids in marine standard reference materials"

### Table of Contents

**Tables**

Table S1: Certified arsenic concentrations (mg/kg) in SRMs 2
Table S2: The instrumental conditions of LC-ICPMS/LC-ESI-QTOFMS 3
Table S3: List of AsLs identified previously and used to develop the PCDL database 4
Table S4: Previously identified arsenolipids in marine biota and seafood SRMs. 9
Table S5: Summary of arsenolipids species identified in SRMs 12
Table S6: The concentration of arsenolipids in standard reference materials 13

**Figures**

Figure S1: Sample preparation and extraction workflow for SRMs 8
Figure S2: The retention time was compared for m/z values of 361.2469, 721.2926, 981.5068, 333.2147, 503.3463, and 749.3240 to the identified product ions for NMIJ CRM 7405-b. 14
Figure S3: The EIC and MS/MS spectra of AsFA502, AsSugPL748, and AsSugPL980, which were identified with lower confidence in NMIJ CRM 7405-b 15
Figure S4: The retention time was compared for m/z values of 475.3163, 361.2450, 333.2130, and 503.3407 with the identified product ions having m/z values of 102.9529, 104.9685, and/or 122.9791 for DOLT-5. 16
Figure S5: The EIC and MS/MS spectra of AsHC332 at m/z 333.2130, and AsHC360 at m/z 361.2450 exhibit fragmentation patterns of 102.9529, 104.9685, and 122.9791 in DOLT-5. 17
Figure S6: The retention time was compared for *m/z* values of 333.2142, 359.2444, and 503.3484 to the identified product ions for BCR-627. 18
Figure S7: The EIC and MS/MS spectra of AsHC358 and AsFA502, which were identified with lower confidence in BCR-627. 19

**References** 20

**Table S1:** Certified arsenic and arsenic species concentrations (mg/kg) in standard reference materials.

| As species | CRM 7405-b[1] | DOLT-5[2] | BCR-627[3] |
|---|---|---|---|
| Inorganic As (V) | 24.4 | - | - |
| Arsenosugar-408 | 1.41 | - | - |
| Arsenosugar-328 | 0.44 | - | - |
| Dimethylarsinic acid | 0.24 | - | 0.15 ± 0.023 |
| Arsenosugar-482 | 0.20 | - | - |
| Arsenosugar-392 | 0.16 | - | - |
| Arsenobetaine | - | - | 3.9 ± 0.2 |
| Total As | 49.5 | 34.6 ± 2.4 | 4.8 ± 0.3 |

**Table S2:** The instrumentation parameters for LC-ICPMS and LC-ESI-QTOFMS.

| Parameter | LC- ICPMS | LC-QTOFMS |
|---|---|---|
| Mode | | Positive |
| Drying Gas Temperature (°C) | | 100 |
| Drying Gas Flowrate (L/min) | | 12 |
| Nebulizer Pressure (psi) | | 45 |
| Sheath Gas Temperature (°C) | | 350 |
| Sheath Gas Flowrate (L/min) | | 11 |
| Nozzle Voltage (V) | | 500 |
| Capillary Voltage (V) | | 4500 |
| Fragmentor Voltage (V) | | 135 |
| Collision Energy (eV) | | 10, 20, and 40 |
| RF power | 1600 W | |
| Spray chamber temperature | 2 ˚C | |
| Sample depth | 8 mm | |
| Mass recorded | *m/z* 75 (As) | |
| Analytical Column | Poroshell 120 EC C18, (4.6×100 mm, 2.7µm, Agilent, USA) | Poroshell 120 EC C18, (4.6×100 mm, 2.7µm, Agilent, USA) |
| Mobile Phase | A. Ultrapure water with 0.05% formic acid<br>B. MeOH with 0.05% formic acid | A. Ultrapure water with 0.05% formic acid<br>B. MeOH with 0.05% formic acid |
| Flow Rate | 0.3 mL/min | 0.3 mL/min |
| Column Temperature | 20ºC | 25ºC |
| Injection Volume | 20 µL | 20 µL |
| Reference Masses | | 121.050873<br>922.009798 |

**Table S3:** List of arsenolipids identified previously and included in the PCDL database.

| Group | Short name | Molecular formula [M] | Expected *m/z* for [M+H]$^+$ (see footnote)$^*$ | Ref |
|---|---|---|---|---|
| Arsenic-containing fatty acids (AsFA) | AsFA220 | $C_7H_{13}AsO_3$ | 221.0151 | 4 |
| | AsFA250 | $C_9H_{19}AsO_3$ | 251.0619 | 4, 5 |
| | AsFA264 | $C_{10}H_{21}AsO_3$ | 265.0779 | 6, 7 |
| | AsFA276 | $C_{11}H_{23}AsO_3$ | 277.0779 | 6, 7 |
| | AsFA278 | $C_{11}H_{25}AsO_3$ | 279.0934 | 4-6 |
| | AsFA292 | $C_{12}H_{25}AsO_3$ | 293.1092 | 5 |
| | thiooxo-AsFA294 | $C_{11}H_{23}AsO_2S$ | 295.0709 | 4 |
| | AsFA302 | $C_{13}H_{23}AsO_3$ | 303.0936 | 6, 7 |
| | AsFA304 | $C_{13}H_{25}AsO_3$ | 305.1092 | 7 |
| | AsFA306 | $C_{13}H_{27}AsO_3$ | 307.1246 | 4, 5 |
| | AsFA316 | $C_{14}H_{25}AsO_3$ | 317.1092 | 6, 7 |
| | AsFA320 | $C_{14}H_{29}AsO_3$ | 321.1405 | 5 |
| | AsFA328 | $C_{15}H_{25}AsO_3$ | 329.1092 | 6, 7 |
| | AsFA334 | $C_{15}H_{31}O_3As$ | 335.1562 | 7 |
| | AsFA342 | $C_{16}H_{27}AsO_3$ | 343.1249 | 6, 8 |
| | AsFA348 | $C_{16}H_{33}O_3As$ | 349.1724 | 9, 10 |
| | thioxo-AsFA350 | $C_{15}H_{31}AsO_2S$ | 351.1325 | 4 |
| | AsFA356 | $C_{17}H_{29}AsO_3$ | 357.1405 | 7 |
| | AsFA360 | $C_{17}H_{33}AsO_3$ | 361.1718 | 11 |
| | AsFA362 | $C_{17}H_{35}O_3As$ | 362.1875 | 6, 7, 12-15 |
| | thio-AsFA362 | $C_{17}H_{35}AsO_2S$ | 379.1651 | 14, 16 |
| | AsFA374 | $C_{18}H_{36}O_3As$ | 375.1884 | 17 |
| | AsFA376 | $C_{18}H_{37}AsO_3$ | 377.2029 | 10 |
| | thio-AsFA378 | $C_{17}H_{35}AsO_2S$ | 379.1635 | 4 |
| | AsFA382 | $C_{19}H_{31}AsO_3$ | 383.1576 | 6 |
| | AsFA384 | $C_{19}H_{33}AsO_3$ | 385.1730 | 11 |
| | AsFA386 | $C_{19}H_{35}AsO_3$ | 387.1875 | 18 |
| | AsFA388 | $C_{19}H_{37}AsO_3$ | 389.2031 | 6, 7, 19, 20 |
| | AsFA390 | $C_{19}H_{39}O_3As$ | 391.2188 | 6, 7, 13 |
| | AsFA404 | $C_{20}H_{41}AsO_3$ | 405.2133 | 7, 13, 14 |
| | AsFA408 | $C_{21}H_{33}AsO_3$ | 409.1718 | 21 |
| | AsFA418 | $C_{21}H_{43}AsO_3$ | 419.2500 | 13, 22 |
| | AsFA422 | $C_{22}H_{35}AsO_3$ | 423.1880 | 6, 13 |
| | AsFA424 | $C_{22}H_{37}AsO_3$ | 425.2032 | 23 |
| | AsFA436 | $C_{23}H_{37}O_3As$ | 437.2031 | 6, 7, 12, 19, 21, 24 |
| | AsFA446 | $C_{23}H_{47}AsO_3$ | 447.2820 | 10 |
| | AsFA448 | $C_{24}H_{37}O_3As$ | 449.2031 | 6, 7, 12, 21, 24 |
| | AsFA462 | $C_{25}H_{39}AsO_3$ | 463.2172 | 6 |
| | AsFA474 | $C_{25}H_{51}O_3As$ | 475.3128 | 10 |
| | AsFA486 | $C_{26}H_{51}O_3As$ | 487.3130 | 10 |
| | AsFA502 | $C_{27}H_{55}O_3As$ | 503.3441 | 10 |
| | AsFA528 | $C_{30}H_{45}O_3As$ | 529.2642 | 25 |
| | AsFA586 | $C_{33}H_{67}O_3As$ | 587.4379 | 14 |

| Group | Short name | Molecular formula [M] | Expected *m/z* for $[M+H]^+$ (see footnote)* | Ref |
|---|---|---|---|---|
| | AsFA612 | $C_{35}H_{69}O_3As$ | 613.4535 | [14] |
| | AsFA732 | $C_{44}H_{81}O_3As$ | 733.5474 | [14] |
| | AsFA746 | $C_{45}H_{83}O_3As$ | 747.5631 | [14] |
| | AsFA760 | $C_{46}H_{85}O_3As$ | 761.5787 | [14] |
| Trimethylarsenio fatty alcohols (TMAsFOH) | TMAsFOH374 | $C_{20}H_{43}AsO$ | 375.2595 | [9, 12] |
| | TMAsFOH418 | $C_{24}H_{39}OAs$ | 419.2284 | [9, 12] |
| | TMAsFOH420 | $C_{19}H_{37}AsO_5$ | 421.1955 | [15] |
| | TMAsFOH436 | $C_{20}H_{41}AsO_5$ | 437.2231 | [15] |
| Arsenic-containing hydrocarbon (AsHC) | AsHC164 | $C_5H_{13}AsO$ | 164.0177 | [26] |
| | AsHC305 | $C_{15}H_{33}AsO$ | 305.1815 | [12] |
| | AsHC330 | $C_{17}H_{35}AsO$ | 331.1969 | [12, 13] |
| | AsHC332 | $C_{17}H_{37}OAs$ | 333.2134 | [12, 19, 22, 24, 27] |
| | thio-AsHC332 | $C_{17}H_{37}AsS$ | 349.1905 | [14, 22] |
| | thio AsHC346 | $C_{18}H_{39}AsS$ | 363.2061 | [22] |
| | AsHC347 | $C_{18}H_{39}AsO$ | 347.2281 | [12, 28] |
| | AsHC358 | $C_{19}H_{39}OAs$ | 359.2281 | [12, 27] |
| | Thio AsHC358 | $C_{19}H_{39}AsS$ | 375.2061 | [22] |
| | AsHC360 | $C_{19}H_{41}OAs$ | 361.2441 | [9, 12, 13, 24] |
| | thio AsHC360 | $C_{19}H_{41}AsS$ | 377.2217 | [22] |
| | AsHC374 | $C_{20}H_{43}OAs$ | 375.2608 | [22, 29] |
| | AsHC 382 | $C_{21}H_{39}AsO$ | 383.2290 | [30] |
| | AsHC386 | $C_{21}H_{43}OAs$ | 387.2602 | [9] |
| | AsHC388 | $C_{21}H_{45}OAs$ | 389.2764 | [28] |
| | AsHC402 | $C_{23}H_{35}AsO$ | 403.1982 | [6] |
| | AsHC404 | $C_{23}H_{37}OAs$ | 405.2134 | [12, 22, 24, 27, 31] |
| | thio AsHC404 | $C_{23}H_{37}AsS$ | 421.1904 | [22] |
| | AsHC406 | $C_{23}H_{39}AsO$ | 407.2295 | [12] |
| | AsHC408 | $C_{23}H_{41}AsO$ | 409.2446 | [32] |
| | AsHC410 | $C_{23}H_{43}AsO$ | 411.2603 | [32] |
| | AsHC440 | $C_{25}H_{49}AsO$ | 441.3077 | [33] |
| | AsHC442 | $C_{25}H_{51}AsO$ | 443.3234 | [33] |
| | AsHC444 | $C_{25}H_{53}AsO$ | 445.3490 | [22] |
| | AsHC542 | $C_{33}H_{55}AsO$ | 543.3547 | [33] |
| | AsHC-OH498 | $C_{25}H_{43}AsO_5$ | 499.2404 | [15] |
| Arsenic-containing phosphatidylcholines (AsPC) | AsPC835 | $C_{41}H_{78}O_9NAsP$ | 836.4780 | [11] |
| | AsPC837 | $C_{41}H_{80}O_9NAsP$ | 838.4940 | [11] |
| | AsPC839 | $C_{41}H_{82}AsNO_9P$ | 840.5090 | [4] |
| | AsPC861 | $C_{43}H_{80}O_9NAsP$ | 862.4930 | [11] |
| | AsPC863 | $C_{43}H_{82}O_9NAsP$ | 864.5080 | [11] |
| | AsPC865 | $C_{43}H_{84}O_9NAsP$ | 866.5240 | [11] |
| | AsPC867 | $C_{43}H_{86}O_9NAsP$ | 868.5400 | [11] |
| | AsPC885 | $C_{45}H_{80}O_9NPAs$ | 886.4950 | [25] |
| | AsPC887 | $C_{45}H_{82}O_9NAsP$ | 888.5080 | [11] |
| | AsPC891 | $C_{45}H_{86}O_9NAsP$ | 892.5400 | [11] |

| Group | Short name | Molecular formula [M] | Expected *m/z* for $[M+H]^+$ (see footnote)* | Ref |
|---|---|---|---|---|
| | AsPC893 | $C_{45}H_{88}O_9NAsP$ | 894.5550 | [11] |
| | AsPC895 | $C_{45}H_{90}O_9NAsP$ | 896.5720 | [11] |
| | AsPC911 | $C_{47}H_{82}O_9NAsP$ | 912.5091 | [25] |
| | AsPC939 | $C_{49}H_{88}O_9NPAs$ | 940.5413 | [9] |
| | AsPC985 | $C_{53}H_{86}O_9NPAs$ | 986.5256 | [9] |
| | AsPC997 | $C_{54}H_{86}O_9NPAs$ | 998.5256 | [9, 25] |
| | AsPL982 | $C_{47}H_{88}O_{14}AsP$ | 983.5205 | [34] |
| | AsPL958 | $C_{45}H_{88}O_{14}AsP$ | 975.4993 | [35] |
| | AsPC911 | $C_{47}H_{82}O_9NAsP$ | 912.5091 | [25] |
| | AsPC939 | $C_{49}H_{86}O_9NAsP$ | 940.5429 | [25] |
| Mono/Di-acyl arsenosugar phospholipids (AsSugPL) | AsSugPL692 | $C_{27}H_{54}AsO_{13}P$ | 693.2591 | [30] |
| | AsSugPL706 | $C_{28}H_{56}AsO_{13}P$ | 707.2748 | [30] |
| | AsSugPL718 | $C_{29}H_{56}AsO_{13}P$ | 719.2747 | [32] |
| | AsSugPL720 | $C_{29}H_{58}AsO_{13}P$ | 721.2904 | [30, 32, 36] |
| | AsSugPL734 | $C_{30}H_{60}AsO_{13}P$ | 735.3060 | [30] |
| | AsSugPL742 | $C_{31}H_{56}AsO_{13}P$ | 743.2747 | [30] |
| | AsSugPL746 | $C_{31}H_{60}AsO_{13}P$ | 747.3060 | [30] |
| | AsSugPL748 | $C_{31}H_{62}AsO_{13}P$ | 749.3217 | [30] |
| | AsSugPL776 | $C_{33}H_{66}AsO_{13}P$ | 777.3530 | [30, 36] |
| | AsSugPL780 | $C_{47}H_{85}AsO_{14}P$ | 981.5043 | [32] |
| | AsSugPL930 | $C_{43}H_{83}O_{14}AsP$ | 931.4897 | [23, 36] |
| | AsSugPL944 | $C_{44}H_{85}O_{14}AsP$ | 945.5049 | [23, 36] |
| | AsSugPL954 | $C_{45}H_{85}AsO_{14}P$ | 955.4890 | [23] |
| | AsSugPL956 | $C_{45}H_{87}O_{14}AsP$ | 957.5018 | [23] |
| | AsSugPL958 | $C_{45}H_{89}O_{14}AsP$ | 959.5205 | [23, 35-37] |
| | AsSugPL972 | $C_{46}H_{91}O_{14}AsP$ | 973.5356 | [23] |
| | AsSugPL978 | $C_{47}H_{84}AsO_{14}P$ | 979.4891 | [32, 37] |
| | AsSugPL980 | $C_{47}H_{86}AsO_{14}P$ | 981.5049 | [23, 36] |
| | AsSugPL982 | $C_{47}H_{88}O_{14}AsP$ | 983.5198 | [23, 32, 36] |
| | AsSugPL984 | $C_{47}H_{90}O_{14}AsP$ | 985.5358 | [23, 36] |
| | AsSugPL986 | $C_{47}H_{92}O_{14}AsP$ | 987.5519 | [23, 34, 36] |
| | AsSugPL998 | $C_{48}H_{90}AsO_{14}P$ | 997.5368 | [36] |
| | AsSugPL1000 | $C_{48}H_{94}O_{14}AsP$ | 1001.5669 | [23, 36] |
| | AsSugPL1006 | $C_{49}H_{88}O_{14}PAs$ | 1007.5200 | [37] |
| | AsSugPL1008 | $C_{49}H_{89}AsO_{14}P$ | 1007.5221 | [37] |
| | AsSugPL1010 | $C_{49}H_{931}AsO_{14}P$ | 1011.5368 | [36] |
| | AsSugPL1012 | $C_{49}H_{95}O_{14}AsP$ | 1013.5681 | [36, 38] |
| | AsSugPL1014 | $C_{49}H_{97}O_{14}AsP$ | 1015.5826 | [35, 36, 38] |
| | AsSugPL1028 | $C_{50}H_{98}O_{14}AsP$ | 1029.5988 | [39] |
| | AsSugPL1040 | $C_{51}H_{97}AsO_{14}P$ | 1040.5910 | [36] |
| | AsSugPL1042 | $C_{51}H_{100}O_{14}AsP$ | 1043.6139 | [28, 35] |
| | AsSugPL1066 | $C_{53}H_{98}AsO_{14}P$ | 1065.5994 | [36] |
| | AsSugPL1070 | $C_{53}H_{104}O_{14}AsP$ | 1071.6452 | [18, 28, 38] |
| Arsenic-containing phosphatidylethanolamines (AsPE) | AsPE819 | $C_{47}H_{77}O_9NAsP$ | 820.447 | [11] |
| | AsPE821 | $C_{47}H_{79}O_9NAsP$ | 822.563 | [11] |
| | AsPE823 | $C_{47}H_{81}O_9NAsP$ | 824.470 | [11] |

| Group | Short name | Molecular formula [M] | Expected *m/z* for [M+H]+ (see footnote)* | Ref |
|---|---|---|---|---|
| | AsPE841 | $C_{49}H_{75}O_9NAsP$ | 842.431 | [11] |
| | AsPE843 | $C_{49}H_{77}O_9NAsP$ | 844.447 | [11] |
| | AsPE845 | $C_{49}H_{79}O_9NAsP$ | 846.463 | [11] |
| | AsPE847 | $C_{49}H_{81}O_9NAsP$ | 848.478 | [11] |
| | AsPE849 | $C_{49}H_{83}O_9NAsP$ | 850.494 | [11] |
| | AsPE1035 | $C_{57}H_{87}O_9NAsP$ | 1036.5387 | [25] |
| Phytyl 2-O-methyl dimethylarsinoyl ribosides (AsSugarPhytol) | AsIsop408 | $C_{18}H_{37}AsO_5$ | 409.1929 | [32] |
| | AsIsop422 | $C_{19}H_{39}AsO_5$ | 423.2086 | [32] |
| | AsIsop464 | $C_{22}H_{55}AsO_5$ | 465.2555 | [32, 40] |
| | AsIsop478 | $C_{23}H_{47}AsO_5$ | 479.2712 | [40] |
| | AsIsop492 | $C_{24}H_{49}AsO_5$ | 493.2868 | [40] |
| | AsIsop506 | $C_{25}H_{51}AsO_5$ | 507.3025 | [40] |
| | AsIsop518 | $C_{26}H_{51}AsO_5$ | 519.3025 | [40] |
| | AsIsop520 | $C_{26}H_{53}AsO_5$ | 521.3181 | [40] |
| | AsIsop532 | $C_{27}H_{53}AsO_5$ | 533.3181 | [40] |
| | AsIsop534 | $C_{27}H_{55}AsO_5$ | 535.3338 | [40] |
| | AsIsop536 | $C_{26}H_{53}AsO_6$ | 537.3131 | [40] |
| | AsIsop546 | $C_{28}H_{55}AsO_5$ | 547.3338 | [37] |
| | AsIsop548 | $C_{28}H_{57}AsO_5$ | 549.3494 | [40] |
| | AsIsop560 | $C_{28}H_{53}AsO_6$ | 561.3131 | [40] |
| | AsIsop562 | $C_{28}H_{55}AsO_6$ | 563.3287 | [40] |

*The preferred ion list for MS/MS analysis mode included the masses for $[M+H]^+$, $[M+Na]^+$, $[M+NH_4]^+$, and their dehydration products, i.e., $[M–H_2O +H]^+$, $[M–H_2O+Na]^+$, and $[M–H_2O+NH_4]^+$

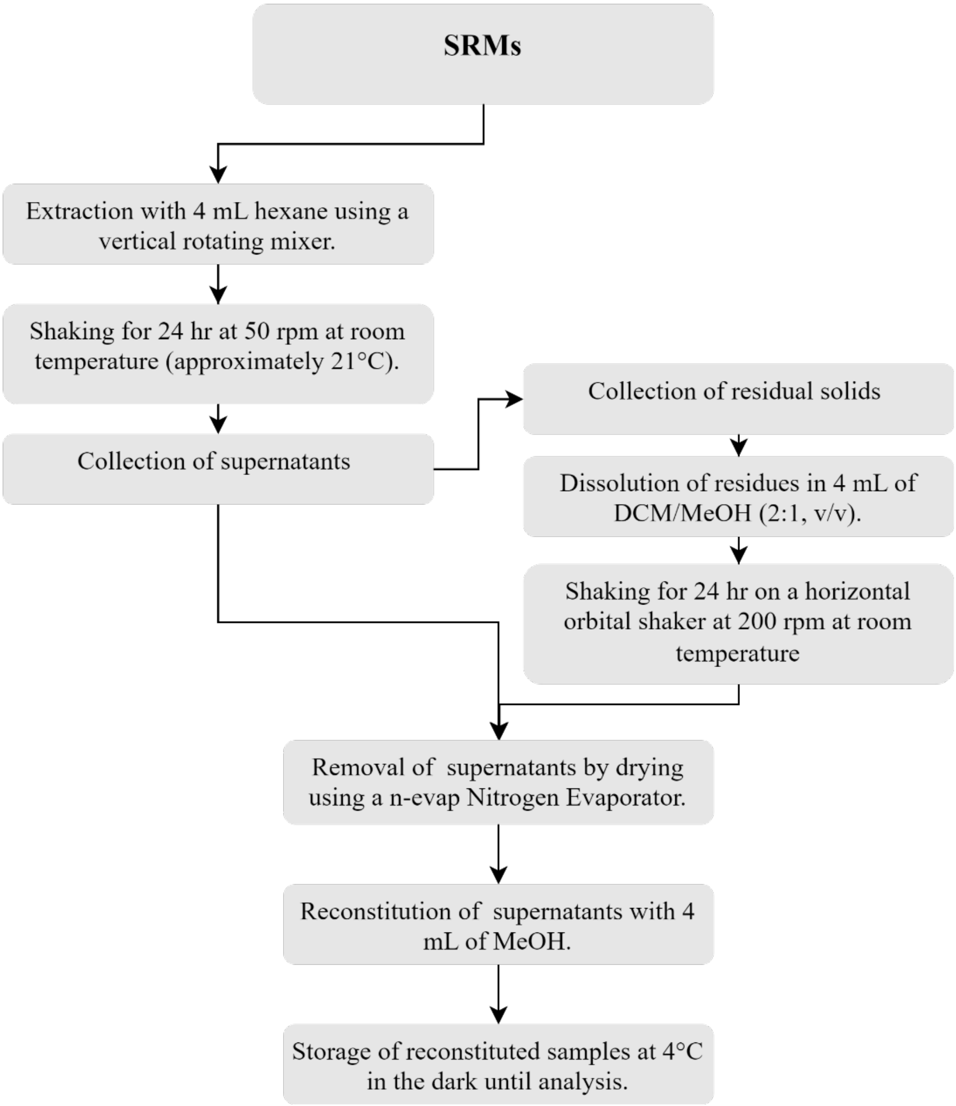


**Figure S1.** Sample preparation and extraction of arsenolipids from standard reference materials.

**Table S4:** Previously reported *m/z* of mass fragments of arsenolipid precursor ions.

| Short name | Molecular formula [$MH^+$] | Precursor ion | Product ions | Reference |
|---|---|---|---|---|
| AsFA264 | $C_{10}H_{22}AsO_3$ | 265.0769 | 247.0684=[M-$H_2$O+H]$^+$<br>122.9786=$(CH_3)_2AsOH_2^+$<br>104.9682=$(CH_3)_2As^+$ | [6, 7] |
| AsFA276 | $C_{11}H_{22}AsO_3$ | 277.0774 | 259.0671= [M-$H_2$O+H] $^+$<br>122.9781=$(CH_3)_2AsOH_2^+$<br>104.9683=$(CH_3)_2As^+$ | [6, 7] |
| AsFA278 | $C_{11}H_{24}AsO_3$ | 279.0934 | 261.0828 = [M-$H_2$O+H]$^+$<br>122.9786 = $(CH_3)_2AsOH_2^+$<br>104.9682 = $(CH_3)_2As^+$<br>1.9 102.9521 = $(CH_2)_2As^+$ | [6] |
| AsFA302 | $C_{13}H_{24}AsO_3$ | 303.0936 | 285.0822= [M-$H_2$O+H] $^+$<br>267.0703=[285.0819- $H_2$O+H] $^+$<br>122.9779=$(CH_3)_2AsOH_2$ $^+$<br>104.9685=$(CH_3)_2As$ $^+$ | [6, 7] |
| AsFA304 | $C_{13}H_{26}AsO_3$ | 305.1094 | 287.0991= [M-$H_2$O+H] $^+$<br>122.9781= $(CH_3)_2AsOH_2^+$<br>104.9685= $(CH_3)_2As^+$ | [7] |
| AsFA316 | $C_{14}H_{26}AsO_3$ | 317.1086 | 299.0989 = [M-$H_2$O+H]$^+$<br>255.1088 = $C_{13}H_{23}As^+$<br>122.9781 = $(CH_3)_2AsOH_2^+$<br>104.9684 = $(CH_3)_2As^+$<br>102.9530 = $(CH_2)_2As^+$ | [6, 7] |
| AsFA328 | $C_{15}H_{26}AsO_3$ | 329.1078 | 311.1008 = [M-$H_2$O+H]$^+$<br>122.9781 = $(CH_3)_2AsOH_2^+$<br>104.9675 = $(CH_3)_2As^+$<br>102.9525 = $(CH_2)_2As^+$ | [6, 7] |
| AsFA342 | $C_{16}H_{28}AsO_3$ | 343.1236 | 325.1141 = [M-$H_2$O+H]$^+$<br>122.9790 = $(CH_3)_2AsOH_2^+$<br>104.9675 = $(CH_3)_2As^+$ | [6] |
| AsFA356 | $C_{17}H_{30}AsO_3$ | 357.1408 | 339.1305 = [M-$H_2$O+H] $^+$<br>122.9790 = $(CH_3)_2AsOH_2^+$<br>104.9683 = $(CH_3)_2As^+$ | [7] |
| AsFA334 | $C_{15}H_{32}O_3As$ | 335.1564 | 317.1457 = [M-$H_2$O+H] $^+$<br>122.9785 = $(CH_3)_2AsOH_2^+$<br>104.9685 = $(CH_3)_2As^+$ | [7] |
| AsFA362 | $C_{17}H_{36}O_3As$ | 363.1875 | 345.1773 = [M-$H_2$O+H]$^+$<br>299.1696 = $C_{16}H_{32}As^+$<br>122.9786 = $(CH_3)_2AsOH_2^+$<br>104.9681 = $(CH_3)_2As^+$<br>102.9525 = $(CH_2)_2As^+$ | [6, 7] |
| AsFA376 | $C_{18}H_{38}AsO_3$ | 377.2029 | 104.9682 = $C_2H_6As^+$<br>122.9763 = $C_2H_7AsO^+$<br>323.1694 = $C_{18}H_{32}As$ -3$H_2$O<br>341.1842 = $C_{18}H_{34}As$ -2$H_2$O<br>359.1931 = $C_{19}H_{38}O_2As$-$H_2$O<br>377.2043 = $C_{18}H_{38}O_3As$ | [10] |

| Short name | Molecular formula [MH$^+$] | Precursor ion | Product ions | Reference |
|---|---|---|---|---|
| AsFA388 | $C_{19}H_{37}AsO_3$ | 389.2036 | 371.1923 = $[M-H_2O+H]^+$<br>122.9786 = $(CH_3)_2AsOH_2^+$<br>104.9683 = $(CH_3)_2As^+$<br>102.9529 = $(CH_2)_2As^+$ | [6, 7] |
| AsFA390 | $C_{19}H_{40}O_3As$ | 391.2195 | 373.2085 = $[M-H_2O+H]^+$<br>122.9788 = $(CH_3)_2AsOH_2^+$<br>104.9678 = $(CH_3)_2As^+$<br>102.9528 = $(CH_2)_2As^+$ | [6, 7] |
| AsFA404 | $C_{20}H_{42}AsO_3$ | 405.2135 | 387.2030 = $[M-H_2O+H]^+$<br>122.9784 = $(CH_3)_2AsOH_2^+$<br>104.9681 = $(CH_3)_2As^+$ | [7] |
| AsFA422 | $C_{22}H_{36}AsO_3$ | 423.1880 | 405.1781 = $[M-H_2O+H]^+$<br>122.9780 = $(CH_3)_2AsOH_2^+$<br>104.9682 = $(CH_3)_2As^+$<br>102.9525 = $(CH_2)_2As^+$ | [6] |
| AsFA436 | $C_{23}H_{38}O_3As$ | 437.2027 | 419.1910 = $[M-H_2O+H]^+$<br>122.9781 = $(CH_3)_2AsOH_2^+$<br>104.9685 = $(CH_3)_2As^+$<br>102.9528 = $(CH_2)_2As^+$ | [6, 7] |
| AsFA448 | $C_{24}H_{38}O_3As$ | 449.2024 | 431.1933 = $[M-H_2O+H]^+$<br>373.1851 = $C_{22}H_{34}As^+$<br>122.9781 = $(CH_3)_2AsOH_2^+$<br>104.9685 = $(CH_3)_2As^+$<br>102.9525 = $(CH_2)_2As^+$ | [6, 7] |
| AsFA462 | $C_{25}H_{40}AsO_3$ | 463.2172 | 445.2050 = $[M-H_2O+H]^+$<br>122.9790 = $(CH_3)_2AsOH_2^+$<br>104.9675 = $(CH_3)_2As^+$<br>102.9528 = $(CH_2)_2As^+$ | [6] |
| AsSugPL720 | $C_{29}H_{58}AsO_{13}P$ | 721.2897 | 97.0289, 237.0101, 329.0573, 409.0236 | [30] |
| AsSugPL748 | | 749.3228 | 97.0240, 237.0103, 329.0574, 409.0242 | [41] |
| AsSugPL776 | $C_{33}H_{66}AsO_{13}P$ | 777.6 | 255.2, 311.3, 391.2, 447.3, 465.3, 483.3, 521.3, 539.3 | [36] |
| AsSugPL930 | $C_{43}H_{85}O_{14}AsP$ | 931.489 | 344.64, 391.01, 409.02, 489.06, 537.54, 700.74, 938.80 | [23] |
| AsSugPL972 | $C_{46}H_{91}O_{14}AsP$ | | 391.011, 437.078 | [23] |
| AsSugPL980 | $C_{47}H_{85}O_{14}AsP$ | 981.505 | 237.010 | [36, 42] |
| AsSugPL984 | $C_{47}H_{91}O_{14}AsP$ | | 451.09, 459.06, and 437.07 | [23] |
| AsSugPL1012 | $C_{49}H_{95}O_{14}AsP$ | 1013.6 | 311.3, 389, 447.3, 463, 481, 539.3, 653.4 | [36] |
| AsPC985 | $C_{53}H_{86}O_9NPAs$ | 986.5250 | 86.0971 = $C_6H_{12}N^+$<br>104.1075 = $C_5H_{14}ON^+$<br>125.0001 = $C_2H_6O_4P^+$<br>184.0735 = $C_5H_{15}O_4PN^+$<br>419.1932 = $C_{23}H_{36}AsO_2^+$<br>437.2025 = $C_{23}H_{36}AsO_3^+$ | [25] |

| Short name | Molecular formula [MH$^+$] | Precursor ion | Product ions | Reference |
|---|---|---|---|---|
| | | | 803.4565 = $C_{48}H_{72}AsO_5^+$ | |
| AsHC330 | $C_{17}H_{35}AsO$ | 331.198 | 104.9689 = $C_2H_6As^+$ | [13] |
| AsHC332 | $C_{17}H_{38}OAs$ | 333.2138 | 104.9682 = $C_2H_6As^+$<br>122.9785 = $C_2H_8AsO^+$<br>175.0455 = $C_7H_{16}As$<br>245.1234 = $C_{12}H_{26}As$<br>315.2002 = $MH^+$-$H_2O$ | [10] |
| AsHC358 | $C_{19}H_{39}OAs$ | 359.2286 | 104.9681 = $C_2H_6As^+$<br>122.9784 = $C_2H_8AsO^+$ | [28] |
| AsHC360 | $C_{19}H_{41}OAs$ | 361.2443 | 104.9681 = $C_2H_6As^+$<br>122.9786 = $C_2H_8AsO^+$<br>343.2345 = $MH^+$-$H_2O$<br><br>104.9678 = $C_2H_6As^+$<br>122.9783 = $C_2H_8AsO^+$<br>343.2336 = $MH^+$-$H_2O$ | [24, 43] |
| AsHC404 | $C_{23}H_{37}OAs$ | 405.2134 | 104.9681 = $C_2H_6As^+$<br>122.9784 = $C_2H_8AsO^+$<br>387.2030 = $MH^+$-$H_2O$ | [43] |

**Table S5:** Summary of the arsenolipid species identified in the hexane and DCM/MeOH fractions of the three standard reference materials investigated.

| SRMs | Hexane | DCM/MeOH | Reference |
|---|---|---|---|
| CRM 7405-b | AsHC360[a] | | [28, 35, 38] |
| | AsSugPL720[a] | AsSugPL720[a] | [28] |
| | AsSugPL980[b] | | This study |
| | | AsSugPL748[b] | This study |
| | | AsFA 502[b] | This study |
| | | AsHC332[a] | [28, 35, 38] |
| DOLT-5 | AsFA474[a] | AsFA474[a] | This study |
| | AsHC360[b] | | [8] |
| | | AsHC332[b] | [8] |
| | | AsFA502[a] | This study |
| BCR-627 | AsHC332[a] | AsHC332[a] | [26, 15, 31] |
| | | AsHC358[b] | [15] |
| | | AsFA 502[b] | This study |

a – high confidence, b – low confidence

**Table S6:** The concentration of arsenolipids in standard reference materials (n=3)

| SRM | Certified Total As (mg/kg) | Total arsenolipids (mg/kg) | Portion of total As present as arsenolipids (%) |
|---|---|---|---|
| NMIJ CRM 7405-b | 49.5 ± 1.0 | 0.50406 ± 0.31136 | 0.5735 - 1.4631 |
| DOLT-5 | 34.6 ± 2.4 | 0.00151 ± 0.00026 | 0.0038 - 0.0049 |
| BCR-627 | 4.8 ± 0.3 | 0.00025 ± 0.00022 | 0.0019 - 0.0091 |

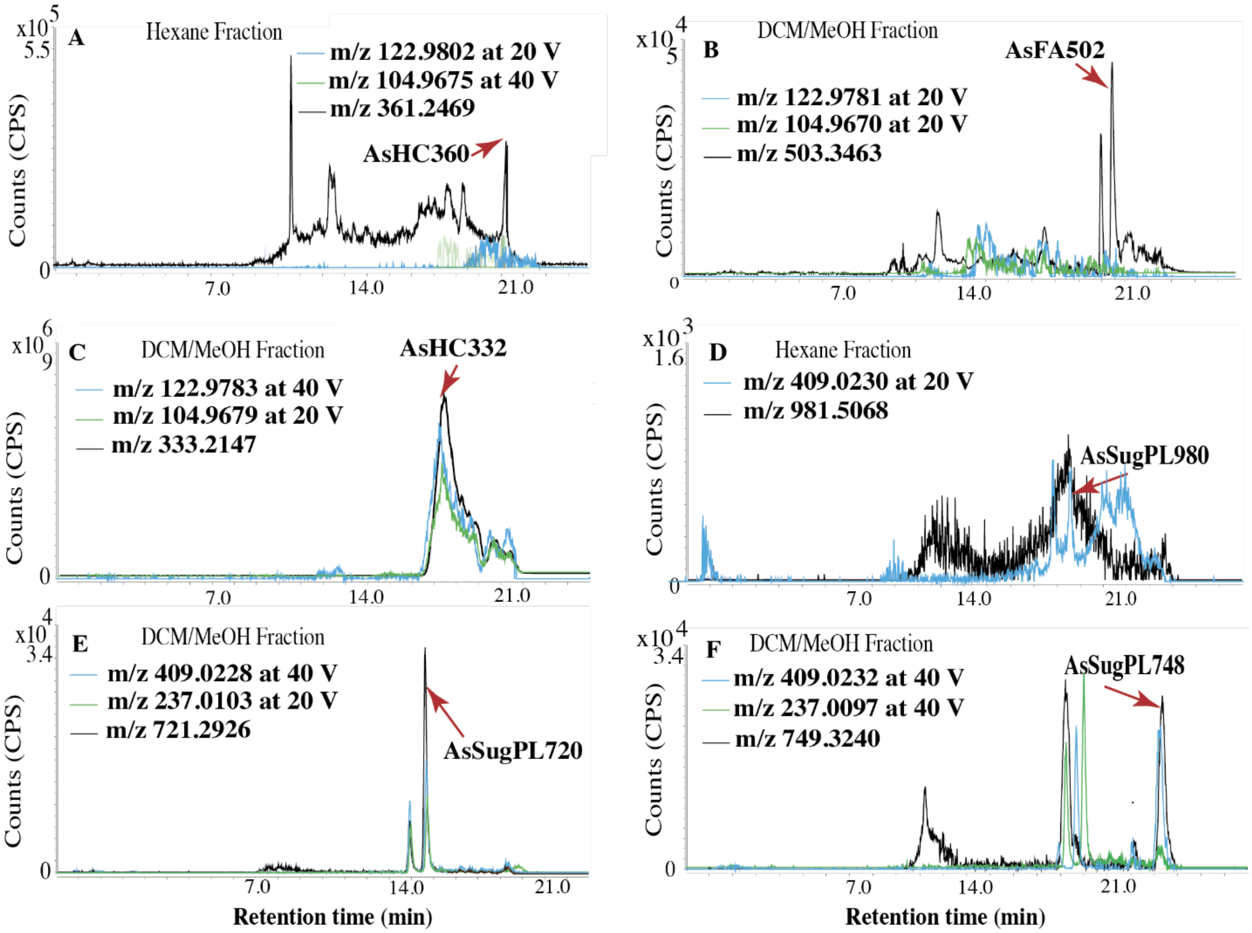


**Figure S2.** Chromatograms of *m/z* values 361.2469 (A), 503.3463 (B), 333.2147 (C), 981.5068 (D), 721.2926 (E), and 749.3240 (F), corresponding to the arsenolipids identified in NMIJ CRM 7405-b: AsHC360, AsFA502, AsHC332, AsSugPL980, AsSugPL720, and AsSugPL748, were overlaid with the chromatograms of *m/z* values of 409.0232, 237.0103, 104.9685, and/or 122.9791, which correspond to the mass fragments associated with these arsenolipid groups. Matching peak retention times verified the presence of the specific arsenolipid compound identified in NMIJ CRM 7405-b. Collision energies (10, 20, or 40 V) at which mass fragmentatio spectra were obtained are indicated in the legend.

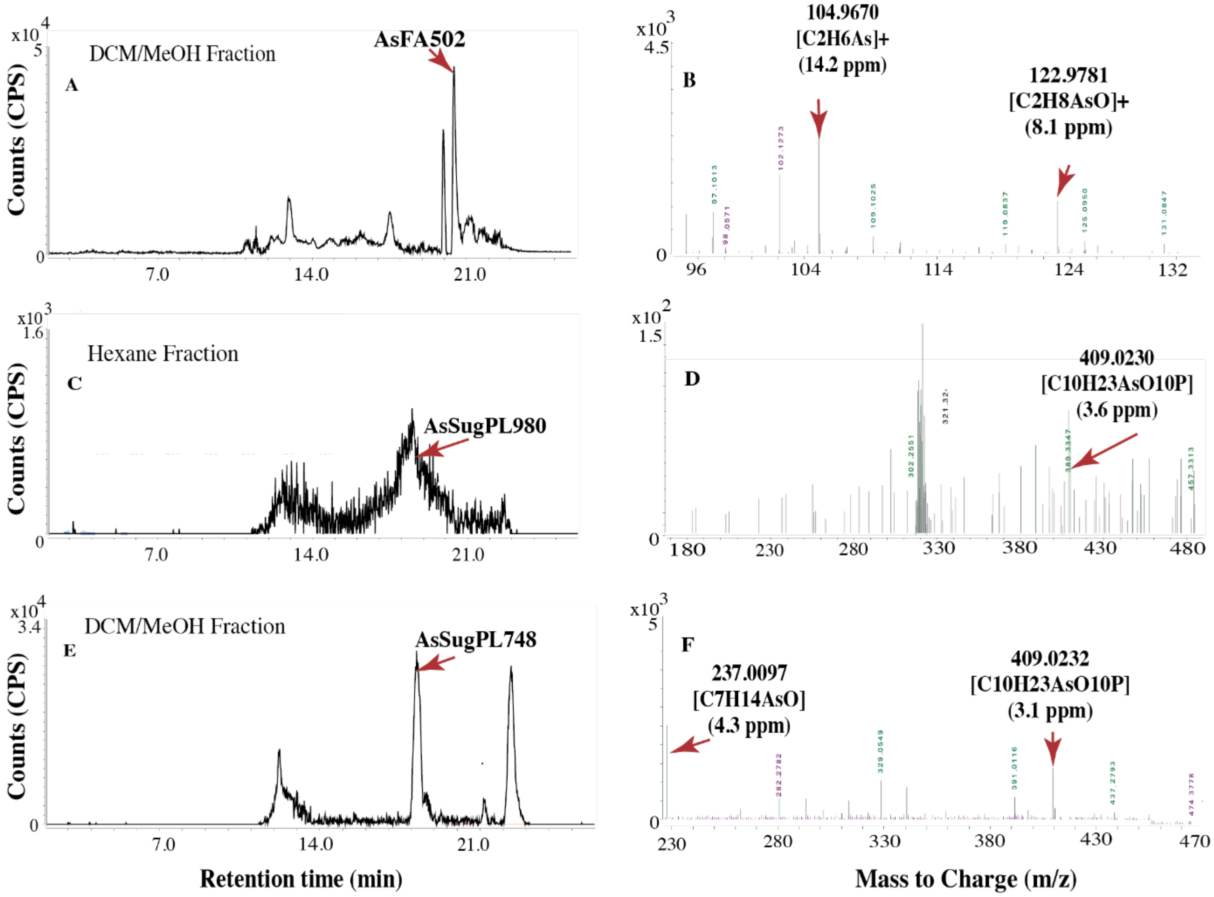


**Figure S3.** The extracted ion chromatograms (EIC) and mass fragmentation spectra of AsFA502 (A & B), AsSugPL980 (C & D), and AsSugPL748 (E & F) in NMIJ CRM 7405-b. Mass fragments with *m/z* 104.9685 and 122.9791 positively identified compounds from the AsFA group, while those with *m/z* 237.0103 and 409.0244 indicated compounds from the AsSugarPL group. The mass error is shown within parentheses.

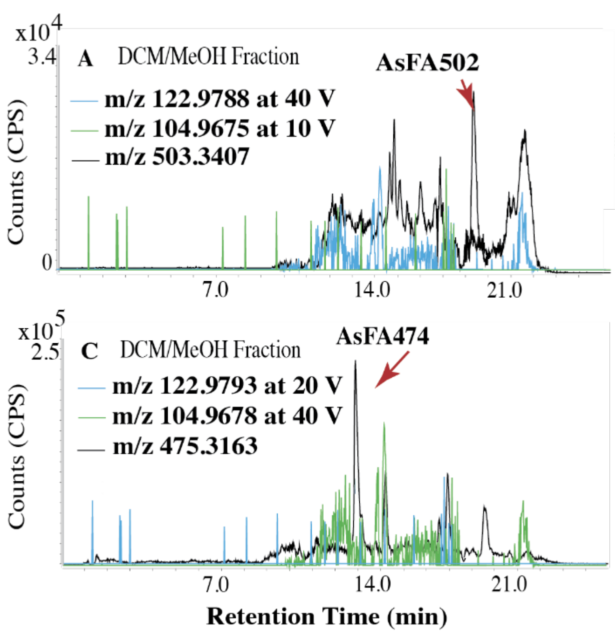


**Figure S4.** Chromatograms of *m/z* values 503.3407 (A), 361.2450 (B), 475.3163 (C), 333.2130 (D), corresponding to the arsenolipids identified in DOLT-5: AsFA502, AsHC360, AsFA474, and AsHC332, were overlaid with the chromatograms of *m/z* values of 102.9529, 104.9685, and/or 122.9791, which correspond to the mass fragments associated with these arsenolipid groups. Matching peak retention times verified the presence of the specific arsenolipid compound  identified in DOLT-5. Collision energies (10, 20, or 40 V) at which mass fragmentatio spectra were obtained are indicated in the legend.

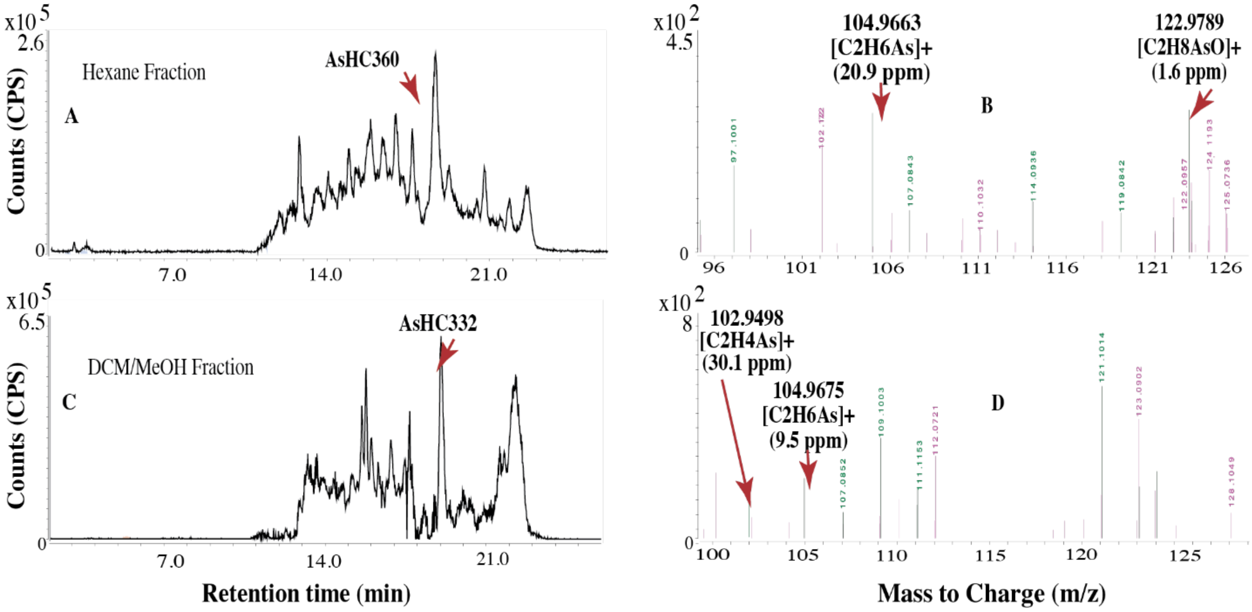


**Figure S5.** The extracted ion chromatograms (EIC) and mass fragmentation spectra of AsHC360 (A & B) at m/z 361.2450 and AsHC332 (C & D) at *m/z* 333.2130 exhibit fragments with *m/z* 102.9529, 104.9685, and 122.9791 in DOLT-5. The mass error calculation is shown within parentheses.

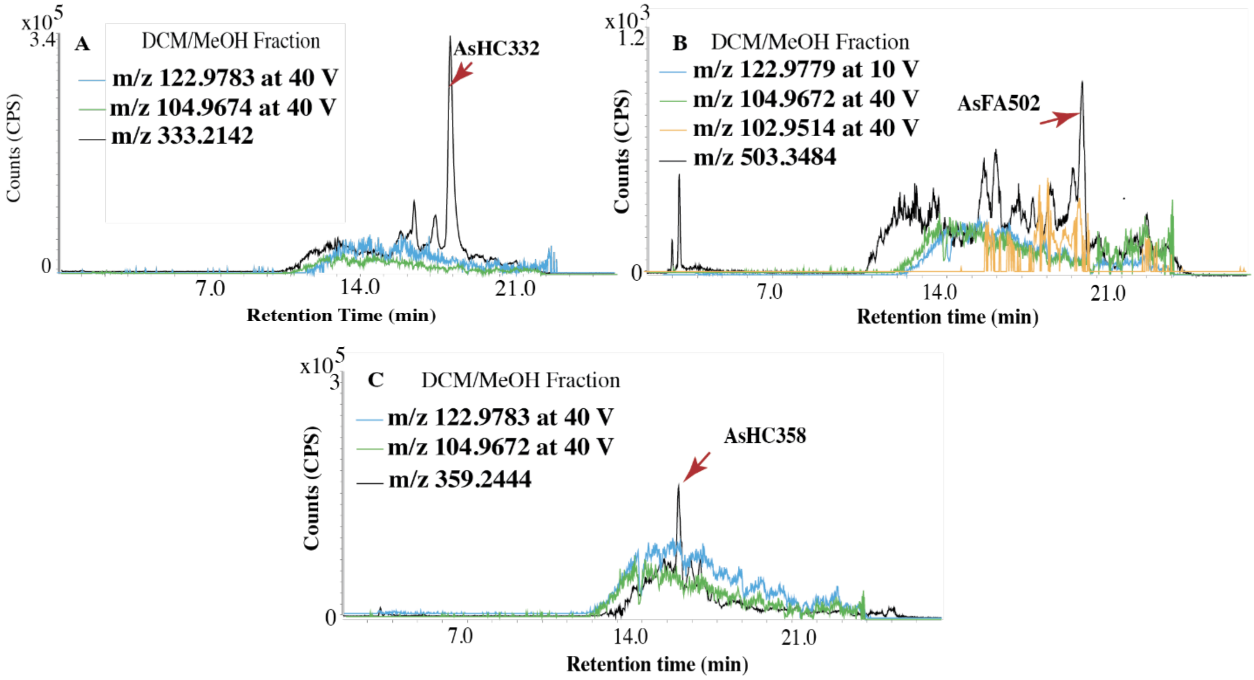


**Figure S6.** Chromatograms of *m/z* values 333.2142 (A), 503.3484 (B), and 359.2444 (C), corresponding to the arsenolipids identified in BCR-627: AsHC332, AsFA502, and AsHC358, were overlaid with the chromatograms of *m/z* values of 102.9514, 104.9672/4, and/or 122.9783/122.9779, which correspond to the mass fragments associated with these arsenolipid groups. Matching peak retention times verified the presence of the specific arsenolipid compound identified in BCR-627. Collision energies (10, 20, or 40 V) at which mass fragmentatio spectra were obtained are indicated in the legend.

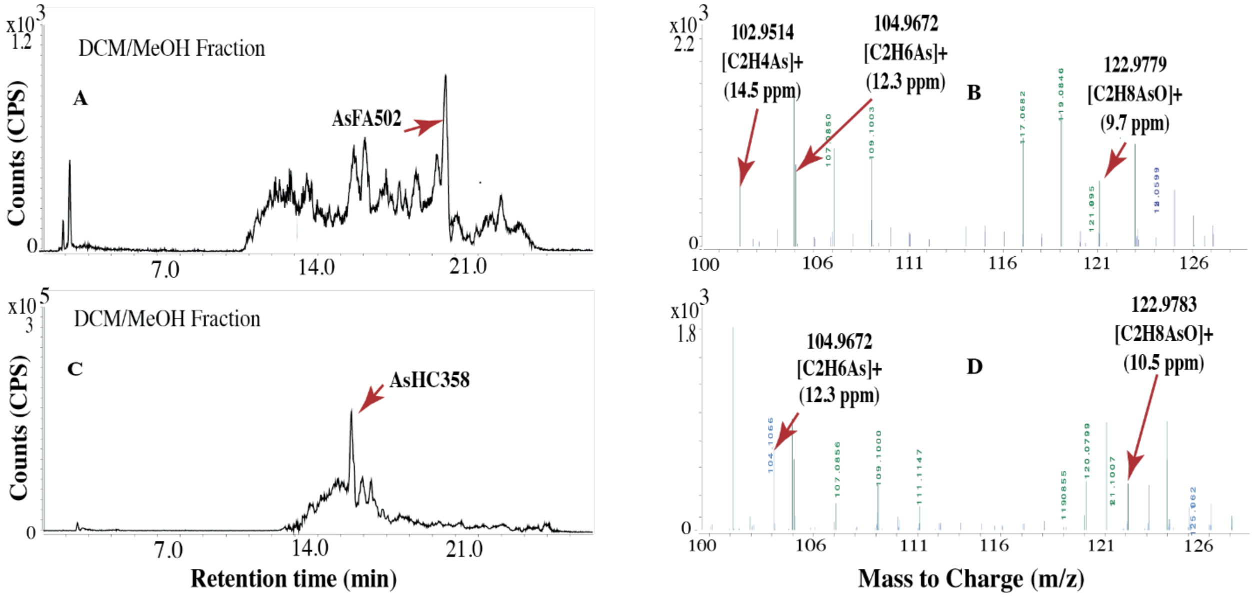


**Figure S7.** Mass fragmentation patterns of AsFA502 (A & B) at *m/z* 503.3484, and AsHC358 (C & D) at *m/z* 359.2623 in BCR-627. Mass error is shown in parentheses.
